# Decoupling Coldness and Softness in Tactile Wetness Perception Using Tunable Hydrogels

**DOI:** 10.1101/2024.09.03.611060

**Authors:** Laura L. Becerra, Nicholas Root, Ainsley Clark, Tarek Rafeedi, William Brown, Alexander X. Chen, Yi Qie, Rachel Blau, Juliana Miller, Kabir Kapadia, Tse Nga Ng, Romke Rouw, Darren J. Lipomi

## Abstract

This study investigates the perception of tactile wetness, a complex sensation experienced by humans. Previous research has primarily focused on either thermal or mechanical cues separately, or has used textiles as stimuli whose parameters are difficult to control. Here, we employed polyacrylamide hydrogels with varying stiffness levels soaked in liquids of distinct thermal conductivities. By psychophysically evaluating participants’ perception of wetness, we showed that the wetness judgments for the samples exhibit a transitive relationship based on the mechanical and thermal cues from an intrinsically tunable organic material. We developed a prediction model of human wetness judgment with an accuracy of 90% and found that the best metrics for the most accurate model were those that were the most human-adjacent: change in temperature at the skin-sample interface (thermal) and compressive force from 2 mm indentation of the sample (mechanical). Given these parameters, we developed a perceptual space capable of recreating 7 distinct levels of wetness perception with the physical parameters used in this study. The results provide insights into the relative contributions of mechanical and thermal stimulus properties in wetness perception. Most notably, this work highlights that the physical characteristics of the skin-stimulus interface can provide ample information for creating a wetness perceptual space, as opposed to the chemical composition of the hydrogels.

## 1 Introduction

Cutaneous wetness is a sensation felt by humans on a daily basis and plays a critical role in behavioral and autonomous adaptation[1]. While this sensation is commonly experienced, there is little attention focused on how the brain perceives this complex sensation on a single scale. The finger has a variety of touch receptors embedded in the epidermal layers. Each class of receptor responds to different types of tactile stimuli depending on its temporal response characteristics, conduction velocity of the nerve fiber, and receptive fields[2, 3]. Stimuli include pressure, temperature, vibration, skin stretching, among others[4]. While there are receptors specific to these stimuli, there is no touch receptor in humans specific to detecting wetness. For this reason it has been hypothesized that wetness is a learned human sensation from tactile experience, triggered by a combination of thermal and mechanical tactile cues[5–7]. Most studies that previously investigated the perception of tactile wetness used stimuli to target one of the two cues (thermal and mechanical)[8–11]. Of the studies that have investigated both thermal and mechanical cues, most have required bulky moving parts to apply pressure for mechanical cues. Or, alternatively, they have used stimuli arising from textiles with variable water content[1]. Thus, a standalone material capable of being altered by intrinsically tuning both mechanical and thermal properties has yet to be tested for tactile wetness perception using the methodologies of psychophysics.

One of the first investigations of moisture perception in humans was described in the Victorian-era work of I.M. Bentley. In this study, the author posed the “blend” hypothesis, which states that wetness perception can be attributed to a combination of temperature and pressure cues[12]. This hypothesis was developed by having participants submerge their finger into a beaker of liquid covered with a thin layer of rubber. Thus, participants were asked to report their sensation of wetness without direct contact with the water. Since Bentley’s blend hypothesis, other works have investigated the roles of mechanical pressure and temperature in wetness perception[1, 8, 9, 13, 14]. Tiest et al. studied static versus dynamic exploration of wet fabrics, where static touch consisted of no mechanical cues, yielding only thermal cues, and dynamic touch included both cues[13]. Their findings concluded that dynamic exploration and, consequently, mechanical cues improve discrimination of wetness in materials compared to only thermal cues. Other studies have focused on the role of surface properties specifically, such as stickiness and friction, on wetness perception[7, 9, 15]. Others have investigated moisture perception relative to levels of sweat production, which psychophysically showed that humans sense skin wetness on their feet primarily from tactile interactions, such as pressure and stickiness, given that thermal cues from skin temperature did not play a role in their wetness perception[16, 17].

Nearly all studies of wetness perception in humans, however, have used textiles with water content as the test material, where participants can come into contact with the water at the interface. To the best of our knowledge after an extensive search, there have only been two studies that looked at temperature and softness (not exerted pressure onto the subject) in wetness perception [18, 19]. The psychophysical results from both studies showed that decreasing temperature of the material led to increased wetness perception in humans. Similar to most of the studies in wetness perception, textiles were used as the test material. To modulate softness, the authors stacked multiple layers of cloth on top of the other, changing overall thickness—and thus stiffness—across samples. One of the studies[19] found that changing thickness of the textiles only resulted in slightly increased softness perception and did not have a significant difference in wetness perception, in disagreement with [18]. Altering softness by stacking samples together can alter heat transfer rates due to inconsistent geometry thickness across samples [20]. To modulate thermal cues, the authors used Peltier elements to directly increase or decrease temperature of the cloth. Although the temperature of the interfaces in these cases was quantified and controllable, the material selection did not offer intrinsic and quantifiable softness nor thermal tunability. Here, we use an organic hydrogel that can be chemically altered to vary inherent softness in a more controlled manner, with similar thicknesses across all samples (1.01 cm +/-0.12 cm) (Fig. 1a,b and Supplementary Fig. 1). Additionally, we control perceived coolness by varying the intrinsic thermal conductivity of the material, as opposed to changing the temperature. Thus, a Peltier element is not required, and inconsistent heat transfer is mitigated with near-identical gel thicknesses across samples. This study, therefore, demonstrates precise control and manipulation of hydrogel composition and corresponding material properties to influence human wetness perception. We use in-depth psychophysics to accurately measure how humans perceive and discriminate between distinct levels of wetness given the varied hydrogel parameters.

**Fig. 1.**
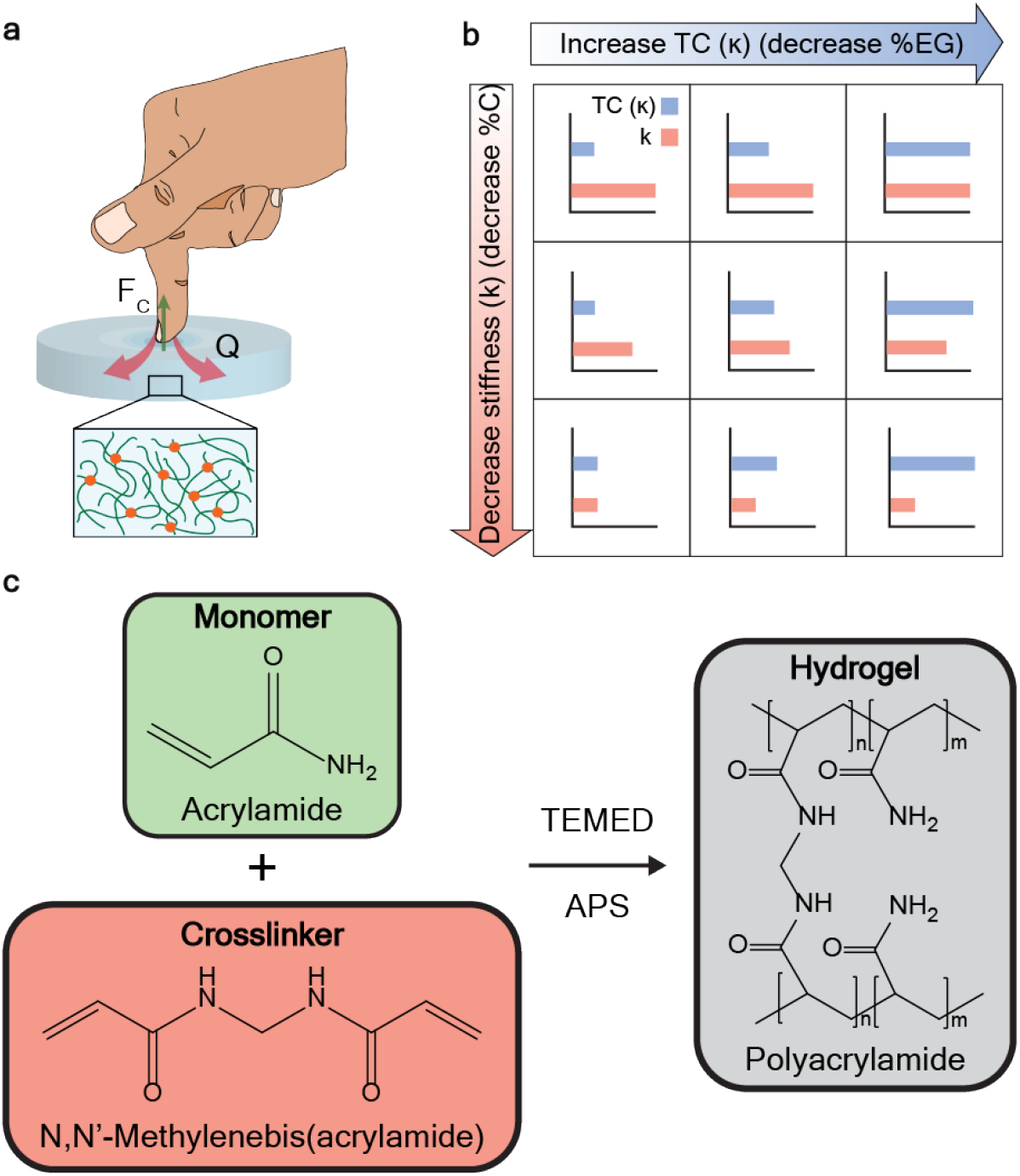
Overview of the experimental methodology. **a**, Schematic diagram of tactile interaction. *F*_*c*_ is the compression force exerted onto the finger from the hydrogel. *Q* is the heat flux across the skin-gel interface. **b**, Matrix of variable properties for each material tested. There are three degrees of thermal conductivity (TC *κ*) in the gel soaking solvents: the increase in TC is achieved by decreasing the percentage of ethylene glycol (EG) relative to water in the gel. There are three degrees of stiffness (*k*), characterized by the ratio of crosslinker-to-monomer ratios. **c**, Chemical structures of the components making up the hydrogels. The polymerization is catalyzed by tetramethlethylenediamine (TEMED) and the initiator is ammonium persulfate (APS).

Polyacrylamide hydrogels are networks of cross-linked acrylamide units which are swollen in the presence of water (Fig. 1c) [21]. Once solidified, these hydrogels are considered biocompatible and safe for handling. These materials have been used for a variety of applications from cell culturing, strain sensing, and drug delivery to the components of flexible batteries[22–24]. Of note for this current study, the mechanical and thermal properties of hydrogels can be easily manipulated. For example, when the fraction of crosslinker relative to the monomer content is increased, the porosity of the gel decreases and stiffness increases[25]. While the liquid component of gels normally consists primarily of water, alternate liquids can be used as well. For example, the viscosity of the liquid can affect the mechanical properties of the gel[26]. We hypothesized that the thermal conductivity of the liquid might also have an effect on the perceived wetness of a gel. Water has a relatively high thermal conductivity compared to most (non-metallic) liquids. Media which are more thermally conductive are perceived as colder than those that are less so, due to their ability to dissipate heat faster from the skin into the medium upon contact[27]. Therefore, hydrogels soaked in liquids with different thermal conductivities should elicit different temperature sensations, at least upon initial contact with the skin. Here, we aimed to evaluate the perception of tactile wetness using gels with varying stiffnesses and containing liquids with a range of thermal conductivities. Our hypothesis was that it would be possible to disentangle the individual contributions of mechanical and thermal cues in the perception of tactile wetness. It should thus be possible to create distinct levels of wetness perception in polymeric solids by modulating the crosslinking density and thermal conductivity.

### 1.1 Touch perception of heat extraction rates

It has been shown that humans are actually better at judging temperature differences rather than absolute temperature[28]. This is due to heat exchange between the skin and object temperature, causing an adaptation to temperature. Adaptation occurs when sensory receptors become less sensitive to a stimulus that was constantly perceived for some time. Detection thresholds for temperature changes are even higher when there are subsequent changes (i.e. heat-cool cycles)[29]. In psychophysics, the increased discrimination abilities with changes in temperature supports using free exploration of physical stimuli as dynamic interactions can yield more information about thermal cues compared to static interactions. Thermal conductivity is a measure of the ability of a material to conduct heat[30]. It plays an important role in thermal contact conductance[31]. To estimate heat extraction across the skin-gel interface, we can use Fourier’s law for calculating heat flow between two bodies in thermal contact, where *Q* is the heat flow (shown in Fig. 1A), *κ* is the thermal conductivity, *A* is the area of the interface, and 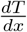 is the temperature gradient across the interface [32] (Eq.1)

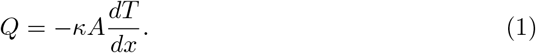

In this study, the heat (*Q*) is traveling from the finger into the gel, which is at ambient temperature in all experiments. Given this direction of flow, we can estimate that the thermal conductivity (*κ*) is dominated by the thermal conductivity of the gel. The interface area is the area of the finger which is in contact with the sample. On average this area has been measured at 2-2.3 *cm*^2^[33].

The change in temperature across the interface can be determined by measuring the interface (or contact) temperature over time upon contact between the finger and the sample. There are two distinguishing characteristics of the temperature curve: the initial rate in temperature drop and the overall change in temperature during the measured period of contact, both of which have been noted as factors influencing the human ability to discriminate temperature [34, 35]. One previous study showed that while thermal cues from initial cooling rates played a prominent role in discriminating materials, thermal cues from total change in temperature also played a role after some time[35]. However, most studies focused on cues from rate of cooling[11, 36, 37]. Interestingly, in this study, we varied thermal conductivity of the gels and show that total change in temperature over 20 seconds to a quasi-steady state temperature resulted in the best prediction model for thermal inputs in wetness perception.

### 1.2 Softness/compliance of materials

Beside perceived temperature, mechanical compliance (“softness”) also influences the perception of wetness. To understand what physical properties influence compliance perception, it is important to identify the measurable characteristics of soft materials. In mechanics of materials, the compliance of a material is the inverse of its stiffness (*k*), which is the ability of a material to resist deformation when applied with a compressive force, shown in Eq. 2, where *F* is the applied force, and *δ* is the displacement of the material caused by the applied force.

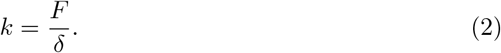

The exploration of haptic perception regarding compliance has undergone extensive study, utilizing stimuli of various stiffness levels[38, 39]. Our group has previously showed that this perception is loosely related to a Hertzian contact model[40]. Metzger et al. revealed that the perception of softness can be influenced by adaptation, suggesting that prior haptic engagements with an object can influence how we perceive the softness of subsequent objects[41]. The susceptibility of adaptation for any sensation suggests that there are neural pathways specialized to detecting different levels of the sensation. Therefore, softness must have distinct neural pathways linked to different softness levels, reinforcing the notion that softness stands as a unique and fundamental aspect of perception. Other studies have highlighted the importance of cutaneous and kinesthetic cues in softness perception[42, 43], and even developed multi-layered perceptual models with high accuracy for softness[44]. A commonly reported conclusion is that surface deformation cues provide the most information for compliance perception[45]. Taking these findings into account, we can incorporate the role of stiffness modulation in stimuli used for wetness perception, and characterize the compression force needed for degrees of surface deformation.

Polyacrylamide hydrogels are considered viscoelastic solid materials[46]. By changing the crosslinker loading in a hydrogel (see Fig. 1b), it has been shown that the elastic modulus, and hence stiffness, of the gel can be tuned[25]. Given the established positive correlation between softness and wetness perception[18], in this study we modulated the softness of hydrogels via different degrees of crosslinking. We characterized differences in compliance via measurements of elastic modulus from shear force, and axial compressive force at a fixed indentation depth.

## 2 Results

### 2.1 Sample design and characterization

#### 2.1.1 Stiffness modulation

The polyacrylamide hydrogel crosslinking densities were varied to create three different levels (0.01, 0.1, and 0.5%), calculated using Equation 3 from a previously published study on tuning polyacrylamide gel stiffness[25].

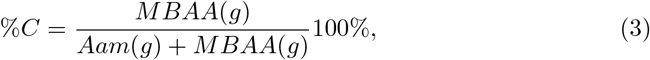

where %C is the percentage of crosslinker in the gel, MBAA(g) is the weight in grams of the crosslinker, N,N’-Methylenebisacrylamide, and Aam(g) is the weight in grams of the monomer, acrylamide, in the gel. After these ratios were combined, a catalyst (TEMED) and initiator (APS) were added to form the gel with deionized water (Fig. 1c).

The resulting porosity both quantitatively (Supplementary Fig. 2) and visually (Fig. 2a) increased with decreasing percentage crosslinker. To quantify how the three solvents used in this study would affect the mechanical properties of the gels, storage modulus and compression force at 2 mm indentation depth (ID) were measured for each of the 9 gels after they were soaked in their solvents. Fig. 2b and c show these parameters as a function of percent crosslinker. Interestingly, compression force at 2 mm indentation depth shows a more consistent trend for each crosslinker amount across soaking solvents compared to storage modulus (i.e. 100% ethylene glycol(EG)) consistently had the highest compressive force and 0% EG had the lowest.) The reason for the discrepancy between mechanical measurement methods could arise from the modes of deformation utilized in the measurements. To measure storage modulus, the gels were placed between two parallel plates with a small amount of axial force and experience shear oscillations at different frequencies. This method can have errors in measurement if the gels are not perfectly flat and flush with the parallel plates. The gels can also slip during oscillations due to residual solvent on the gel even after drying the outside. A movement that is more similar to the way a subject’s finger would interact with a gel is compressive force upon indentation. This method also mitigates potential errors from slippage and the greater need for flush contact in rheological measurements, providing a more applicable physical variable to correlate with the psychophysical responses from participants. When plotting these two measurements against each other (Fig. 2d), we observe a much lower modulus of the 0.5 %C gel with EG compared to the other two solvents, almost matching the modulus of the 0.1 %C gel with EG.

**Fig. 2.**
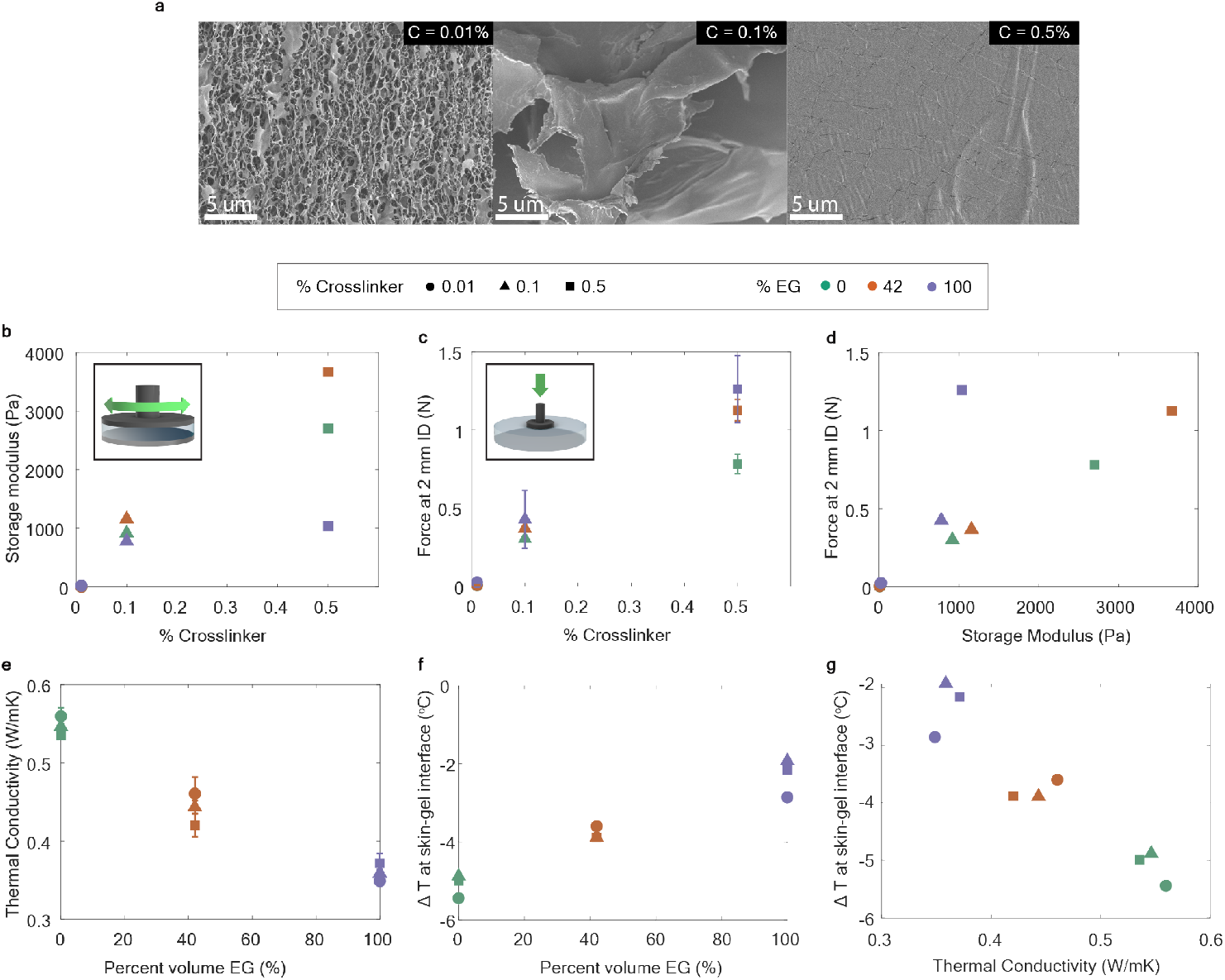
Material characterization. **a**, SEM images of hydrogels of each crosslinking density (increasing crosslinking density from left to right). **b**, Storage modulus calculated from shear mechanical force across crosslinker content. **c**, The compression force from indenting the gel at 2 mm across crosslinker content. Center values are averages of the measurements on a different location for each sample and error bars were calculated using the corresponding standard deviation (n=3). **d**, Compression force at 2 mm ID as a function of storage modulus. **e**, Thermal conductivity of hydrogels as a function of EG content. Center values are averages of multiple measurements for each sample and error bars were calculated using the corresponding standard deviation (n=3).**f**, Change in temperature at the gel/skin interface upon stimulus contact as a function of EG content. **g**, Change in temperature at gel/skin interface as a function of thermal conductivity across all gels.

#### 2.1.2 Thermal conductivity modulation

The EG dilutions chosen as the soaking solvents for the gels were selected based on a pilot experiment using 0.1%C hydrogels soaked in 5 different dilutions of glycerol. From a series of initial pilot psychophysical experiments (two-alternative forced choice (2-AFC)), a minimum TC difference of 0.125 W/mK that was detectable 50% of the time was measured at the corresponding abscissa and used as the minimum difference needed in the gel soaking solvents (Supplementary Fig. 3). As we observed that glycerol content significantly affected the stiffness of the gel, we instead chose to use EG for all future experiments. Using thermal conductivity values from literature, three dilutions of EG in water were used to soak one gel of each crosslinker amount: 0, 42, and 100 vol% EG. These reported mixture thermal conductivities are 0.595, 0.429, and 0.251 W/mK, respectively[47], satisfying the threshold determined in the pilot experiment.

**Fig. 3.**
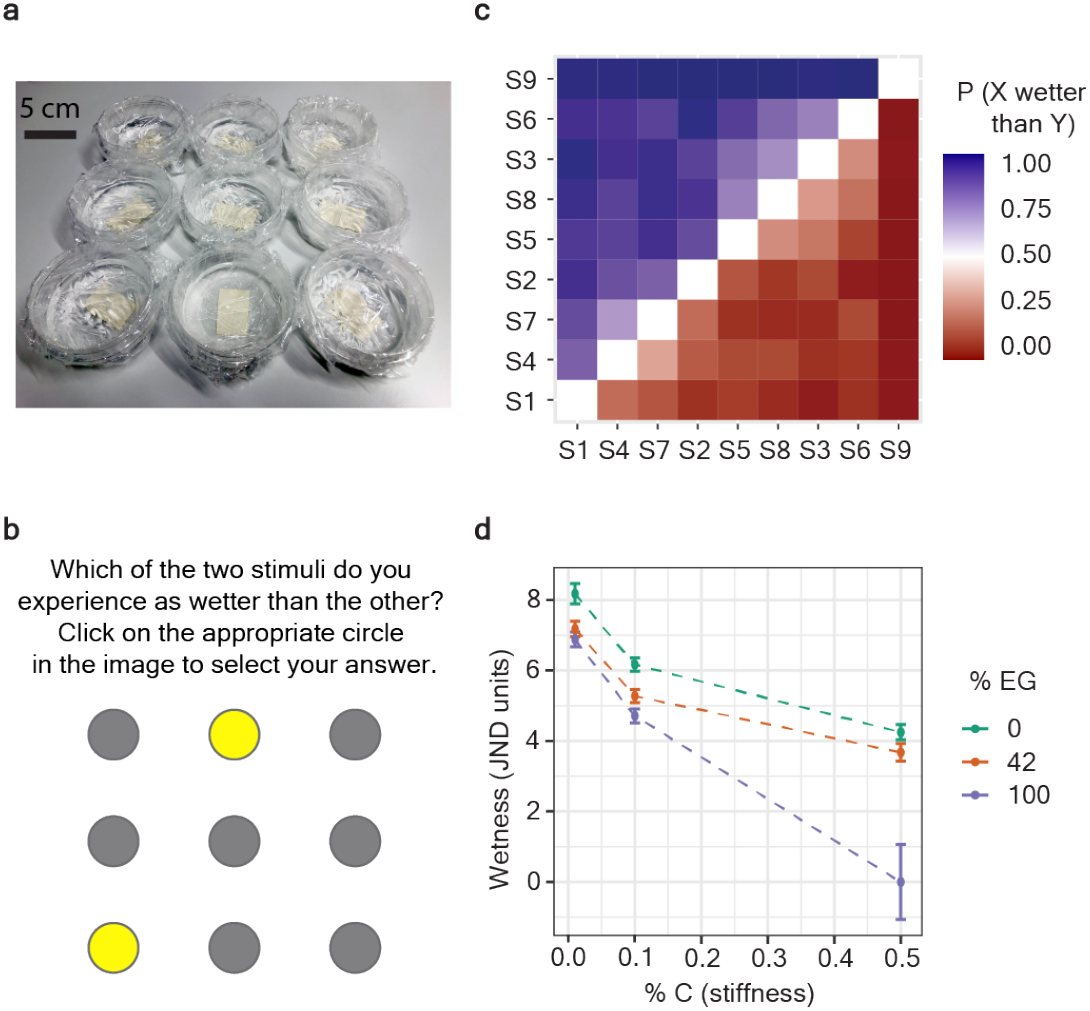
Psychophysical results. **a**, Photo of gels in psychophysical test setup **b**, Graphical user interface for comparison instructions and stimulus selection **c**, Resulting transitivity matrix of all pairwise comparisons of stimuli **d**, Relationship between perceived wetness, and stimulus composition (percent crosslinker and percent EG) in the paired comparison psychophysics experiment (N=20 participants). Y-axis values are Bradley-Terry ability score parameters normalized to JND scale (divided by log(3)). Error bars are 95% confidence intervals on the parameter estimates.

Nonetheless, we characterized the thermal conductivity of the gels soaked in the mixtures across all degrees of crosslinking (Fig. 2e). The overall trend showed a non-overlapping distribution in thermal conductivities across EG content. Generally, a higher volumetric fraction of EG in the soaking solution produced lower thermal conductivity gels. This trend is consistent with the reported thermal conductivities of standard EG-water mixtures without gels [47]. Similar to the characterization techniques in the softness modulation, we sought to measure a thermal attribute more adjacent to the touch interaction felt by the user. Hence, the skin/gel thermal interaction was captured by fitting the characteristic cooling curve of the interface upon contact. We measured the temperature at the skin/gel interface over 20 s upon initial contact (Supplementary Fig. 4) and applied a log fit regression line for psychophysical analysis and fit an exponential (decay) regression to estimate the rate of temperature change and final temperature (Supplementary Fig. 5). Figure 2f and 2g show the total temperature change at the interface for each gel and thermal conductivity as a function of %EG, respectively.

**Fig. 4.**
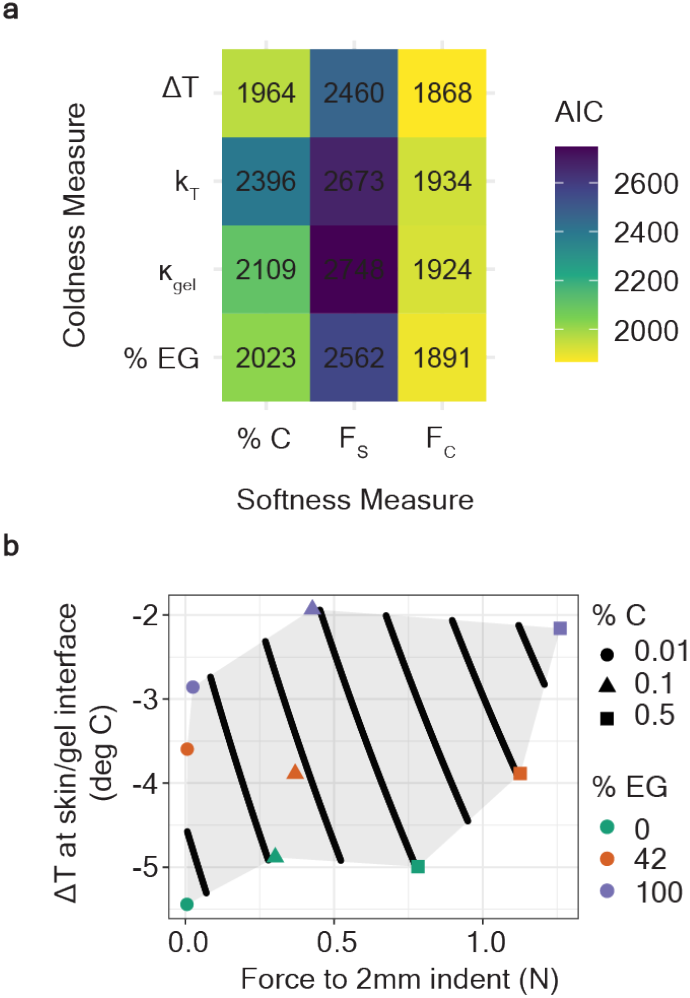
Psychophysical results. **a**, Prediction model comparison (AIC values) across possible input parameters for coldness (Δ*T* is total change in interface temperature, *k*_*T*_ is the rate of change of interface temperature, *κ*_*gel*_ is the gel thermal conductivity, %EG is percent EG in the soaking solvent) and softness measures (%C is percent crosslinker of the gel, *F*_*s*_ is shear force, *F*_*c*_ is compression force) **b**, JND space from human-like physical properties with markers for stimuli recipe composition (%C and %EG) overlaid. The black lines represent distinct levels of wetness perception.

As crosslinker content, and therefore stiffness, decreased, the change in temperature at the skin-gel interface also decreased across all solvents. The increase in contact area at the interface with the softer gels allows for more thermal pathways with a higher contact area (*A*), increasing overall heat flux (*Q*) from the gel to the finger. Additionally, it has been psychophysically shown that temperature changes can be better resolved if the area of thermal stimulation increases [34]. Moreover, experimental measurements and molecular dynamic simulations have revealed that the thermal conductivity of PAam hydrogels is influenced by their crosslinking density, as demonstrated by Tang et al. in their study [48]. In particular, they examined a range of cross-linker concentrations (including those used in this study) and measured the thermal conductivities. The authors observed a rising trend in thermal conductivity for gels with crosslinker concentrations ranging from 0.016 to 0.099 mol%, which is likely attributed to the increase in thermal pathways resulting from the transition from van der Waals interactions to covalent bonds. However, at higher crosslinker concentrations (0.099-0.263 mol%), a decrease in thermal conductivity was noted, and this was attributed to an escalation in phonon scattering arising from an increase in side chains. In our results, we saw a general trend of lower crosslinking density correlating to higher thermal conductivity and vice versa, even for the gel that fell into the crosslinking range that should decrease in thermal conductivity according to Tang et al. The only exception to the trend was when 100% EG was used as the soaking solvent, causing the opposite effect (decreasing crosslinking density = decreasing thermal conductivity). The high EG content likely causes a different interaction between the gel matrix and the surrounding medium, potentially reducing gaps in the matrix where solvent would be, resulting in a more dense, thermal insulating material.

### 2.2 Psychophysical studies

The nine gels in the study were presented to each subject as shown in Fig. 3a. Each gel in the matrix was covered with a thin layer of polymer film. Participants were instructed to compare two of the gels in each trial (2-AFC) and to select which of the two gels they experienced as being wetter using a computer interface (Fig. 3b).

Fig. 3c depicts all pairwise comparisons: the percentage of the time (across all participants) that Sample X was perceived as wetter than Sample Y. The stimuli have been arranged on the X and Y axis such that the left-(bottom-) most stimulus was perceived overall (across all pairwise comparisons) to be the wettest stimulus, and the right-(top-) most stimulus was perceived overall to be the least wet stimulus. Crucially, this arrangement yields a plot in which all pairwise comparisons below the diagonal are p *<* 0.5, and all pairwise comparisons above the diagonal are p *>* 0.5. This property means that the data satisfies the criterion of weak stochastic transitivity: if Sample A is perceived as wetter than Sample B and Sample B is perceived as wetter than Sample C, then Sample A is also perceived as wetter than Sample C. Transitivity of wetness judgements is important because it implies that even though two different physical parameters were manipulated in our study, participants still experienced wetness on a unidimensional scale [49].

To more precisely characterize participants’ perceptions, we used a Bradley-Terry model [50] to transform pairwise comparison data into a latent “wetness” variable that reflects the relative probability that a stimulus will be judged as wetter than another. Bradley-Terry parameters are in log odds units; therefore, dividing all strength estimates by log(3) yields a wetness scale in units of Just Noticable Difference (JND), such that when Stimulus A is one unit wetter than Stimulus B, it will be perceived as wetter 75 percent of the time. Figure 3d depicts the perceived wetness levels of each of the stimuli in our study, as a function of how much crosslinker-to-monomer content was used for each gel (x axis), and how much EG-to-water content there was in the soaking liquid (graph color).

Qualitatively, it appears from Figure 3d that both stiffness and thermal conductivity influence human perceptual judgements of wetness. To quantify this observation, we fit a series of Bradley-Terry models in which the wetness of each stimulus depended (via a linear predictor) [51] on each possible combination of our measurements of stiffness (%C, *F*_*s*_, *F*_*c*_) and coldness (Δ*T, k*_*T*_, *κ*_*gel*_, %EG) and their interaction. In all 12 models, stiffness and coldness were highly significant predictors of the pairwise comparison results (Likelihood Ratio Tests, Bonferroni correction for multiple comparisons, all p *<* 0.001).

Each model can be compared using the Akaike Information Criterion (AIC), which quantifies the relative prediction error of each model (Figure 4a). Lower values of AIC indicate that one model is more predictive than other model; for example, the odds that the best model (AIC = 1868) is better than the second-best model (AIC = 1891) are 98716 to 1 (calculated as *e*^(Δ*AIC/*2)^); an AIC difference of 10 is typically sufficient to establish that one model is significantly more predictive [52]. The strongest pattern in the AIC values was in choice of stiffness measurement: across all choices of coldness measurement, models using force to 2 mm indent as the measurement of stiffness were significantly better than models using crosslinker ratio or storage modulus. Furthermore, across all choices of stiffness measure, models using total change in skin temperature were significantly better than models using rate of change of skin temperature, thermal conductivity, or percent EG.

We next compared actual human judgements of wetness to the predictions of the best-fitting model (force to 2mm indent as the measurement of stiffness, total change in skin temperature as the measure of coldness). Receiver Operating Characteristics (ROC) analysis indicates that the model performed significantly better than chance, with an accuracy of 90.0% and an area under the curve (AUC) of 0.94 (bootstrapped 95% CI = [0.93,0.95]). To more rigorously test our model, and to estimate its out-of-sample generalizability, we re-fit the model on subsets of data in which a stimulus was removed, and tested the model’s accuracy at predicting the stimulus it had not seen “Leave one out cross validation”, LOOCV). When predicting human judgements for an unseen stimulus, the LOOCV model still performed significantly better than chance, with an accuracy of 79.8% and an AUC of 0.87 (bootstrapped 95% CI = [0.84,0.90]).

Finally, we used the parameters of the best-fitting model to generate predictions of human judgements across the gamut of possible physical parameters we can generate. This yielded a representation of perceptual space that can be used to recreate a particular level of perceived wetness using different combinations of physical stimulus parameters (Fig. 4b). The diagonal lines in Fig. 4b depict combinations of parameters yielding the same perceived wetness. For example, the sample with 0.1%C and 100%EG is perceived to be approximately as wet as the sample with 0.5%C and 0%EG, but these samples are perceived to be 1 JND drier (i.e., judged to be drier 75% of the time) than the sample with 0.1%C and 42%EG. Notice that there are at most 7 JND steps that fit within the perceptual space that can generated by the combination of physical parameters that our technique can produce. This means that the techniques developed in this study are capable of developing stimuli with 7 different levels of wetness that can be reliably distinguished from each other.

## 3 Methods

### 3.1 Hydrogel synthesis

The hydrogels were synthesized using the following protocol. First, aqueous stock solutions were prepared in water for the synthesis, shown in Table 1.

**Table 1.** Stock solution concentrations for hydrogel synthesis.

| Solution | Concentration |
| --- | --- |
| Acrylamide (Aam) | 0.5 g/mL |
| N,N'-Methylenebisacrylamide (MBAA) | 0.025 g/mL |
| Ammonium persulfate (APS) | 10 %wt/vol |

Each gel consisted of 20.00 mL (7.034 M) acrylamide stock solution, 250 *µ*L (0.438 M) of APS stock solution, and 25 *µ*L of TEMED. The volume of MBAA stock solution (0.1622 M) used for each crosslinker amount is as follows: 0.04 mL (C = 0.01%), 0.2 mL (C = 0.05%), 0.4 mL (C = 0.1%), and 2 mL (C = 0.5%). The volume of water needed for the solution to reach a total volume of 50 mL per gel was added. Acrylamide, MBAA, and APS stock solutions were added to the appropriate volume of water in a 150 mL Erlenmeyer flask. The flask was swirled gently to promote mixing, then the TEMED was added. After another gentle swirl, the solution mixture was transferred to a petri dish. The petri dish was then placed under UV light (350 nm) to cure the gels. Once cured, the gels were placed in water to allow for gel expansion. After the gels had swelled, the water was replaced twice, once each subsequent day. On the third day, they were transferred to one of three solvent mixtures: pure water, 42% EG by volume, or pure EG. To ensure proper absorption of the solvent, the new solvents were switched out three times over the following days.

The gels were cut into circles using a circle-shaped cookie cutter with diameters of 6 cm and had thicknesses of *∼*1 cm. For the experiment, stimuli were presented in nine circle shaped polyethylene containers (diameter = 10 cm, height = 5 cm), each containing a polyacrylamide hydrogel submerged in the soaking solvent. A thin film of bisphenol Afree plastic wrap was placed over the gels in the containers. The leak-proof containers were labelled from S1 to S9 in a way that was not visible to the participants.

### 3.2 Hydrogel characterization

#### 3.2.1 Rheology

A rheometer (TA Instrument Discover HR-30 Hybrid) was used to measure the storage modulus of all hydrogels. Each gel was cut to have a 38 - 40 mm diameter to match the area of the sandblasted Peltier parallel plate geometry. The outer layer of the gel was dried with a delicate task wipe to avoid slippage during measurements. Each gel underwent an amplitude sweep at 25^*°*^C with an angular frequency of 1 rad/s.

#### 3.2.2 Force-indentation measurements

Mechanical properties of the gels were further investigated using compressive force at 2 mm indentation depth. A motorized compression test stand (Mark-10 ESM303) was used with a force gauge to measure the compression force from 0 mm to 2 mm indentation at a rate of *∼*0.65 mm/min. The compression attachment making contact with the gels was a flat disc with a diameter of 12.7 mm. The samples were pat dry and the soaking solvents were removed from the containers for measurement to avoid slippage upon compression.

#### 3.2.3 Porosity

To measure the porosity percentage of each gel crosslinker amount, small gel samples were cut and lyophilized (Labconco FreeZone freeze-dryer 2.5L, 115V, 60Hz) for 48 h. Each freeze-dried sample was weighed and then soaked in absolute ethanol for 48 hours. After soaking, the volume and weight of each sample were measured. The percentage porosity of each sample was then calculated using Equation 1, where *M*_1_ and *M*_2_ are the mass of the hydrogel sample before and after soaking in absolute ethanol, *ρ* is the density of the absolute ethanol, and V is the volume of the hydrogel after swelling.

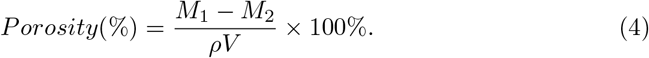

#### 3.2.4 Scanning electron microscopy

One gel for each crosslinker amount (0.01, 0.05, 0.1, and 0.5%) was imaged via scanning electron microscopy (Zeiss Sigma 500). The gels were prepared by being placed on dry ice for 3 h until frozen and immediately transferred to lyophilizer (Labconco FreeZone freeze-dryer 2.5L, 115V, 60Hz) and were freeze-dried for 48 h. The gels were then sputtered with Iridium using a sputter coater (Emitech K575X) with 70% power intensity for 60 sec. After sputtering, the gels were imaged.

#### 3.2.5 Temperature at gel-skin interface

The temperature profile at the gel-skin interface was measured using two K-type thermocouple probes connected to a USB temperature data logger (LASCAR ELUSB-TP-LCD). The probe was stripped so that the sensing portion of the wire was exposed and taped to the fingertip where contact would be made with the gel. The other probe tip was placed on the gel itself. Each measurement consisted of 10 s of baseline data before contact, followed by 30 s of temperature data from contact with the gel. The finger was submerged in the film covered samples until the gel was felt with light force.

#### 3.2.6 Thermal conductivity measurements

Thermal conductivity of each gel was measured using a thermal properties analyzer (METER TEMPOS) with a 6 cm single needle probe (KS-3). Each gel had a volume of 100 mL and was cured in a custom designed 3D-printed mold. The mold had an inner diameter of 42 mm and a height of 96 mm, and was designed specifically to meet the minimum radial distance of the medium around the needle probe for accurate measurements. The gels were all initially soaked in water for 3 days, with the water switched out everyday. On the fourth day, the gels were then soaked in their respective EG-water mixtures for 3 days, with the solutions switched out each day. For a single measurement, the gel was removed from the solution and placed in a cylindrical glass dish to hold the gel in place. The probe was connected to the meter and the needle was fully inserted vertically into the top center of the gel. The measurements were done inside of a Styrofoam box to mitigate large heat fluctuations. Three measurements were taken per gel.

### 3.3 Psychophysical experimental design

#### 3.3.1 Participants

20 participants (13 female; 18-50 years old; mean age = 24.71) were recruited from the University of Amsterdam. Participants volunteered (n=13) or received course credit (n=7). The experiments in this study were approved by an Ethics Review Board at the University of Amsterdam (ERB ID 2023-BC-15923). The methods were carried out in accordance to the ERB’s guidelines and regulations. Informed consent was obtained from all participants in the study.

#### 3.3.2 Procedure

Before participants arrived, the experimenter arranged the nine experimental stimuli in a 3×3 grid in front of the experiment computer (Fig. 3a). The placement of each stimulus within the grid was randomized (by computer algorithm) for each participant. Upon arrival, participants used a sink in the experiment room to wash and dry their hands (to avoid fouling the experimental materials, and to reduce variations in skin moisture). They were then seated in front of the experimental setup, read and signed the consent form, and read the instructions for the experimental task.

The experimental task was a 2-AFC paired comparison procedure in which the difference in perceived wetness of each pair of stimuli was evaluated. On each trial, participants were instructed to touch two of the nine stimuli simultaneously, using their left and right index fingers. Their task was to indicate which stimulus felt wettest to them by selecting the corresponding circle on a computer screen (Fig. 3b). There was no time constraint: participants could take as long as they wished to make each judgement. There were a total of 144 trials, divided into four blocks of 36 trials, one for each of the 36 possible pairwise comparisons (i.e., each pairwise comparison was judged a total of four times). Within each block, the trials were presented in randomized order. Between each block, participants were required to take a break of at least 30 seconds.

## 4 Discussion

In this study, we psychophysically tested human perception of wetness using an organic material with tunable mechanical and thermal properties. Our aim was to decouple the influences of coldness and softness on tactile wetness perception and to understand which aspects of the stimulus dominate in wetness judgment. Change in coldness was enabled by soaking the hydrogels in liquids of different thermal conductivities, changing the heat extraction rates from the finger into the gel. Softness was varied by changing the stiffness of the gel via number of crosslinks in the polymer matrix.

Our psychophysical analysis revealed a consistent trend—softer and colder properties elicited an enhanced perception of wetness, aligning with findings from prior studies[5, 8, 14, 53]. That is, decreased stiffness of the material and rapid dissipation of heat from the finger correlated with an increased perception of wetness. Notably, participants checked their fingers after experiment completion and reported being surprised their fingers were dry. Emphatic reporting of wetness even in the absence of direct liquid contact, echoed observations made by I.M. Bentley [12]. We found a predictable pattern in the participants’ judgment of wetness by showing the transitivity across the different combinations of stiffness and coldness, implying that the perceptual relationships of these two parameters are not arbitrary.

Further analysis revealed that parameters directly measuring the mechanical and thermal experiences of the subjects (such as changes in temperature at the interface and compression force at a 2 mm indentation depth) yielded the most effective prediction model (AIC). These results are notable for at least two reasons. First, the most widely-accepted “gold standard” measurements of stiffness (storage modulus) and thermal conductivity were amongst the poorest predictors of wetness perception. Second, the best predictors of human perception were in some sense the most “human-adjacent” measurements, less closely related to intrinsic properties of the stimuli and more related to how they interact with a finger-like object when pressed. Additionally, the total change in interface temperature was a better predictor in coldness measure for wetness perception than the rate of change of interface temperature. This is interesting because a common trend in thermal perception literature claims that while the amplitude of temperature change is a factor in temperature perception, cooling rates deem more important[35]. While thermal perception may rely heavily on initial cooling rates in temperature discrimination, our model suggests that perhaps long-term thermal characteristics are more indicative of wetness, such as prolonged sensation of cooling after contact due to evaporative cooling on wet skin.

Given that wetness is a more complex sensation compared to temperature, humans may require more stimuli exploration time to judge wetness than needed to judge temperature. Filingeri et al. concluded that while skin cooling influences wetness perception, other sensing modalities, such as vision, can affect individual thresholds for complex sensations such as wetness [11]. Therefore, it is possible that the modulation of softness affected the temperature discrimination abilities in this study. Lastly, our temperature measurements, taken using a thermocouple taped to the finger, likely yielded more stable estimates of the asymptotic temperature. Since the final temperatures were recorded after collecting numerous data points to a quasi-steady state, these measurements are less susceptible to the noise typically associated with the rapid changes observed in initial cooling rates.

We leveraged the prediction model to predict combinations of change in interface temperature and compression force that yield approximately equal sensations of wetness (in Fig 4b, stimuli on the same black line), and combinations that yield sensations of wetness that are approximately one JND apart (in Fig 4b, stimuli on adjacent black lines). Given these material types and stimulus parameters, we can expect to perceive approximately 7 gradations of wetness levels. The ability to correlate hydrogel synthesis parameters/recipes with the 7 perception levels is useful, but the more impactful result, is the ability to correlate it to the interface stiffness and temperature profile. This will enable a much more broad application of wetness stimuli, not restricting it to parameters specific to hydrogels.

This study represents a significant advancement towards developing a versatile, all-in-one material for rendering technology. By inherently tuning both thermal and mechanical properties, this material effectively targets specific wetness sensations, offering multiple benefits. Thermally, the material’s thermal conductivity eliminates the need for external temperature adjustments, allowing it to function optimally at room temperature. Mechanically, the material maintains a consistent macroscopic geometry while still enabling stiffness modulation, enhancing its utility and simplifying its application. In essence, our findings contribute both to the field of psychophysics and also hold implications for material design in human-machine interfaces, highlighting the intricate interplay between softness and coldness in shaping the perceptive experience of wetness.

## Supplementary information

All supplementary information is provided in an appended document.

## Acknowledgments

L.L.B. acknowledges the support provided by the National Science Foundation Graduate Research Fellowship Program under Grant DGE-2038238, the Achievement Rewards for College Scientists (ARCS) Foundation, and the UC President’s Dissertation Year Fellowship at UC San Diego. N.R. and R.R. acknowledge support from the Dutch Research Council (NWO) under Grant 406.21.GO.021.

R.B. acknowledges funding from the European Union’s Horizon 2020 Research and Innovation Programme under the Marie Sklodowska-Curie grant agreement No 898571.

## Declarations

### Ethics approval

The experiments in this study were approved by an Ethics Review Board at the University of Amsterdam (ERB ID 2023-BC-15923).

### Consent to participate

Participants provided informed consent to participate in this study.

### Availability of data and materials

The data that support the findings of this study are available from the corresponding author upon reasonable request.

### Code availability

All code used for psychophysical statistical analysis is available upon request.

### Authors’ contributions

L.L.B. conceived the study’s idea and material selection and modification. L.L.B., A.C., and W.B. synthesized study samples. L.L.B., T.R., W.B., A.X.C., Y.Q., R.B. and J.M. contributed to sample characterization experiments. N.R. and R.R. designed, carried out, and analyzed all psychophysical experiments of the study. D.J.L. and R.R. oversaw the study. T.N.N. and N.R. provided guidance for figure design and layout. All authors contributed to the writing of the manuscript.

