## Supplementary Information for "Decoupling Coldness and Softness in Tactile Wetness Perception Using Tunable Hydrogels"

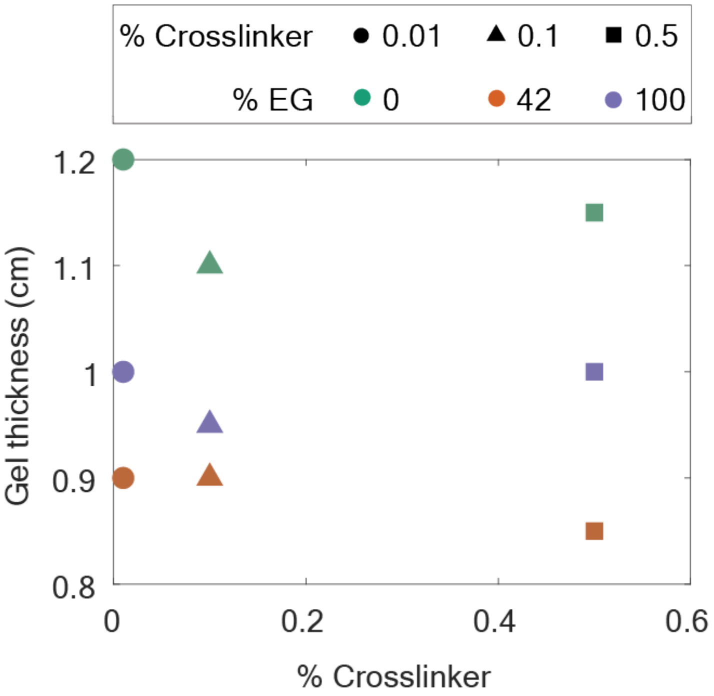


**Supplementary Fig. 1 Gel thickness measurements** NOTE: Legend information is taken from Figure 2 of the main text.

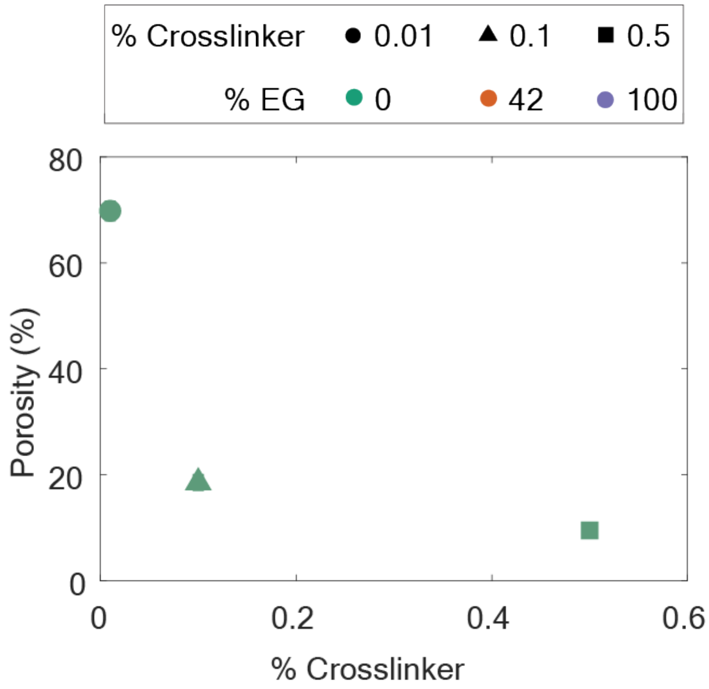


**Supplementary Fig. 2 Gel porosity** NOTE: Legend information is taken from Figure 2 of the main text.

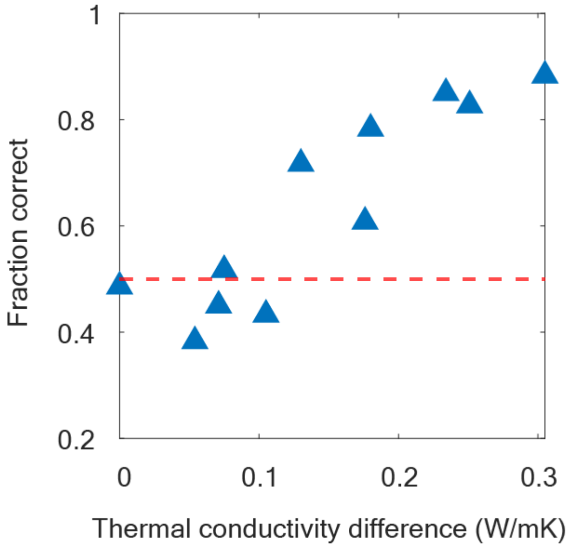


**Supplementary Fig. 3 Pilot experiment for thermal conductivity threshold**

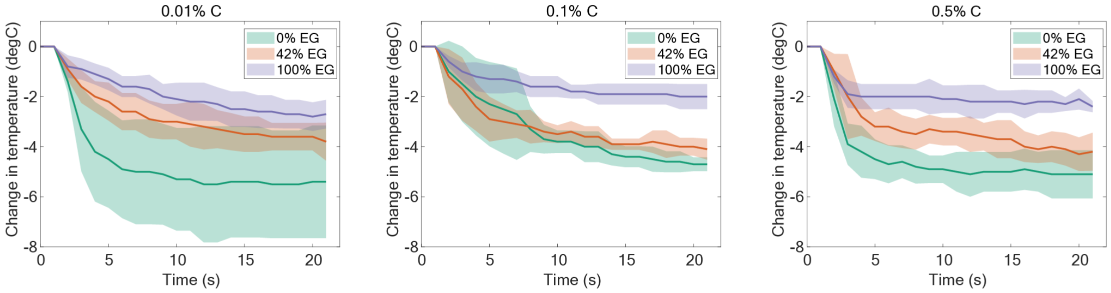


**Supplementary Fig. 4  Skin-gel temperature curves**

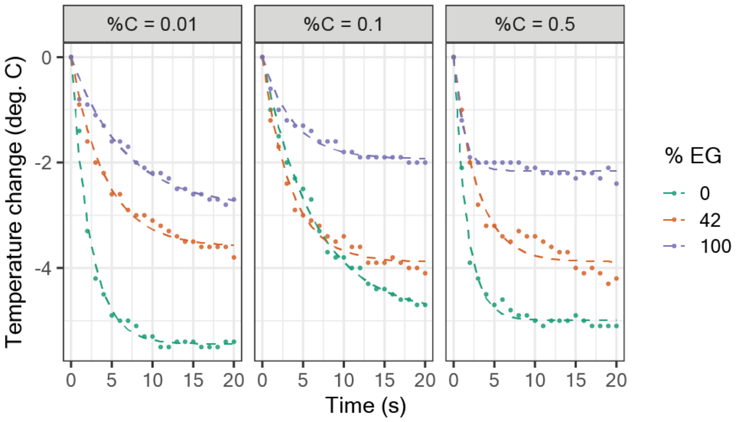


**Supplementary Fig. 5  Function-fitted skin-gel temperature curves**
